# In vitro model for resistance in oncogene-dependent tumors at the limit of radiological detectability

**DOI:** 10.1101/756593

**Authors:** Nina Müller, Johannes Brägelmann, Carina Lorenz, Ulrich P. Michel, Dennis Plenker, Sandra Ortiz-Cuaran, Jonathan Weiss, Reinhard Büttner, Martin Peifer, Roman K. Thomas, Martin L. Sos, Johannes Berg

## Abstract

In solid tumors, the response to targeted therapy is typically short-lived, as therapy-resistant mutants can quickly expand during therapy. Here we analyze the spectrum of such resistance mutations coexisting in a large population of cancer cells. We use an iterative scheme of artificial evolution to amplify and isolate different resistance mechanisms. As a proof of concept, we apply our scheme to PC-9 cells, a human non-small cell lung cancer cell line with an activating EGFR mutation. The mechanisms we find comprise the well-known gatekeeper-mutation T790M in EGFR, a mutation in NRAS, the amplification of MET-ligand *HGF*, as well as induction of AKT-mTOR signaling. In this model, a combination of four drugs targeting these mechanisms prevents not only the expansion of resistant cells, but also inhibits the growth of drug-tolerant cells, which can otherwise act as a reservoir for further resistance mutations. These data suggest that a finite number of drugs specifically acting on individual resistant clones may be able to control resistance in oncogenically driven lung cancer.

## 1 Introduction

In a sufficiently large population of tumor cells, resistant cells exist already prior to targeted therapy [1, 2, 3, 4]. This is because a single point mutation in a specific genomic position can make a cancer cell resistant to a particular targeted drug. While the rate at which such mutations arise is small, the large number of cells in a solid tumor can lead to a substantial number of resistant mutants: At a diameter of 1 cm, a size at which early detection is possible using radiological imaging [5], a solid tumor contains about 10^8^ cells [6]. A mutation rate of 10^−8^–10^−9^ mutations per genomic site and cell division [7] then yields an expected number of 0.1–1 point mutations at a specific site arising at every generation. These estimates for the mutation rate and the number of cancer cells are conservative, so the total number of mutants arising at every generation will typically be higher than this. Hence, at the time treatment begins, a solid tumor typically contains cells carrying a resistance mutation. Under treatment, cells that are sensitive to therapy are eradicated, while resistant cells expand. Eventually, resistant cells repopulate the tumor, leading to the acquired resistance phenotype [8].

In this paper, we use a cell-line model to study the resistance mutations existing in a population of cancer cells corresponding in size to a tumor at the threshold of radiological detectability. We use populations of up to 5 × 10^8^ cells, which exceeds the pools containing a few millions of cells used in previous studies [3, 4, 9]. To amplify and isolate resistant mutants, we establish an iterative scheme of artificial evolution experiments. In this way, low-frequency mutants which are too rare to be detected by direct sequencing can be studied. This allows to address the spectrum of resistance mechanisms found in a population of cancer cells: How many distinct mechanisms are present at the beginning of therapy and what are their biochemical bases? What are their growth rates? Can they all be targeted by extant compounds? These questions need to be answered to design polytherapeutic approaches where drugs are combined upfront or switched between in order to suppress the emergence of resistant cancer cells [10, 11, 12].

We apply our approach to PC-9 cells, a human non-small cell lung cancer cell line with an activating mutation in the epidermal growth factor receptor (EGFR). Tumors driven by an activating mutation in EGFR frequently respond to the EGFR tyrosine kinase inhibitor erlotinib [13, 14]. Mutations conferring resistance to erlotinib treatment include the well-studied EGFR gatekeeper mutation T790M [15, 16], copy number alterations in *MET* and *HER2*, but also mutations that activate bypass signaling at the level of MAPK and PI3K signaling, see [17, 11, 18] for reviews on resistance mechanisms and [19] for a review of EGFR signalling. A similar picture holds for targeted therapies acting on other mechanisms like *ALK*-rearranged tumors, where several ALK mutations, as well as increased EGFR signaling and *KIT* amplification have been identified as possible resistance mechanisms [20, 21, 22, 17]. As the different resistance mutations arise independently, mutant cells with different resistance mechanisms can coexist in a large population of tumor cells, leading to polyclonal resistance [23, 24, 3, 12]: The different resistance mechanisms compete with each other once non-resistant cells have been eliminated by therapy (Fig. 1A).

**Figure 1:**
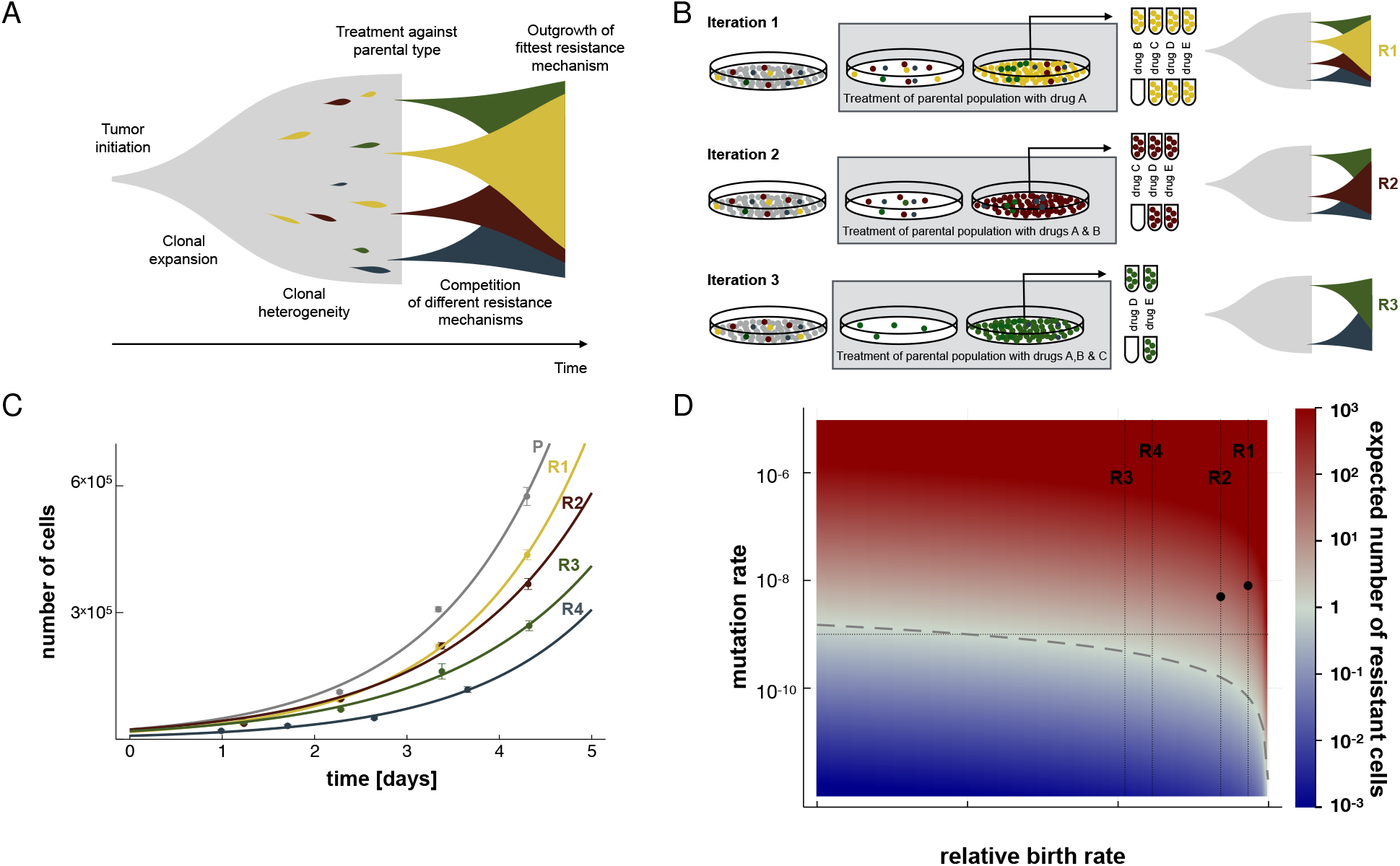
Resistant mutants in a large population of sensitive cells. A The schematic plot show the population dynamics of treatment-sensitive cells (gray) and treatment-resistant cells carrying different resistance mutations (red, yellow, green, and blue) both before and during treatment. The sensitive cells are eradicated under treatment, while the resistant cells expand and compete with each other, eventually leading to a tumor with acquired resistance. B Each step of our iterative scheme starts with a large population of 5 × 10^8^ cells and uses compounds eradicating the resistant cells identified in previous rounds, see text. C The resistant cell lines R1–R4 isolated in this manner grow at different rates, as seen by tracking their population size over a four-day period without treatment. Solid lines are exponential fits. D The expected number of resistant mutants cells increases as a function of their birth rate (relative to the parental cells) and their mutation rate. Red hues indicate an expected number of resistant cells higher than one, indicating parameters where resistant cells typically exist in the population prior to therapy. Vertical lines correspond to the measured growth rates for R1–R4, black dots indicate estimates of point mutation rates, see text and SI B. The horizontal line indicates base-line rate of point mutations of 10^−9^ per nucleotide and cell division.

Our key finding is that, for PC-9 cells, a set of four drugs is sufficient to suppress all resistance mechanisms present in a large population. The drugs in this combination are specific to the different resistance mechanisms, unlike the combinations of antibiotics or anti-retroviral drugs used to control resistance in chronic bacterial or viral infections. Interestingly, this set of compounds also prevents the expansion of drug tolerant persisters, which otherwise provide an alternative route to resistance [25, 4, 9]. These drug-tolerant resisters can also be eliminated, for instance by inhibiting the anti-apoptotic Bcl-2 [4]. Eliminating all resistance mechanisms, as well as the drug-tolerant persisters clears a large population of PC-9 cells: neither the outgrowth of colonies nor individual surviving cells are observed after withdrawing the treatment.

## 2 Results

Our starting point is a population of approximately 5 × 10^8^ PC-9 cells. This number roughly corresponds to a tumor of size 1cm^3^ [6]. Treatment with 500 nM of the EFGR inhibitor erlotinib quickly eradicates cells susceptible to erlotinib, leaving cells carrying one of the resistance mutations, see SI A for a sample time-course and dosing. As we show below, cells with different resistance mutations grow at different rates, so over time the resistance mechanism with the highest growth rate will outcompete all other mechanisms. The resulting cells (termed R1) are harvested and screened against a combination of erlotinib and compounds from a panel of 17 inhibitors (Fig 1B and SI A). This panel of compounds was chosen to target a wide range of known and potential resistance mechanisms. We identify the compound the cell line R1 is most susceptible to (together with erlotinib) and then iterate our approach: We return again to a large population of parental PC-9 cells and treat with erlotinib and the compound identified in the previous step. By construction, this eliminates not only all cells susceptible to erlotinib, but also eradicates R1, leaving the remaining mechanisms to compete with one another. The fastest growing mutant (termed R2) outcompetes all others and is again harvested for screening and analysis.

In principle, each iteration of this scheme selects a further resistance mechanism and identifies compounds that the newly derived resistant cell line is sensitive to. In each step, the bulk of susceptible cells is rapidly depleted under high drug doses (several times the IC_50_ concentration, see SI A), while resistant cells existing in the population expand. Our scheme thus differs from inducing a resistance mechanism via long-term exposure of a small population to a low drug dose [26, 1], or letting the population pass through a drug-tolerant persister state [25, 4, 9]. Each iteration takes about 4 to 6 months depending on the growth rate of the resistant cells.

### Cells with different resistance mechanisms have different growth rates

Based on the above experimental scheme, we successively isolated four different erlotinib-resistant cell lines labelled R1-R4. We used cell counting at different time points to measure the growth rates *g* of the different resistant lines in the absence of therapeutic compounds. R1 grows at 96% percent of the rate of the parental PC-9 cells, R2 at 91%, R3 at 74% and R4 at 79% (corresponding to growth rates *g*_*P*_ = 0.82 0.01, *g*_*R1*_ = 0.79 0.01, *g*_*R2*_ = 0.75 0.01, *g*_*R3*_ = 0.61 0.02 and *g*_*R4*_ = 0.65 0.01 per day, respectively, see Fig. 1C). The resistant cell lines thus grow more slowly than the parental cells, as is to be expected since otherwise they would take over the population.

The growth rate affects how many cells carrying a given resistance mechanism exist in a population of a given size. Using a simple population genetic model, we determine the expected number of mutant cells as a function of growth rate, mutation rate, and population size (Fig. 1D and SI B and C). Using an order-of-magnitude estimate for the somatic point mutation rate of 10^−9^ per generation [7], we can use the measured growth rates to estimate the number of cells with a given resistance mechanism present in the population prior to therapy. Our model predicts an average of approximately 120 R1 cells in a population of 5 × 10^8^ cells, or about one cell in 4 × 10^6^. Similarly, we expect about 30 R2 cells in a population of 5 × 10^8^. The expected number of resistant cells decreases when their growth rate decreases. For the slowest-growing cell line, R3, a mutation rate of 10^−9^ per generation corresponds to only 2 resistant cells on average in 5 × 10^8^. Resistance mechanisms with substantially lower growth rates would thus typically be absent from a population of this size. In these estimates, we adjusted the mutation rates for the size of the mutational targets, see below and SI D. In the case of R1, which turns out to carry the EGFR T790M mutation, the eight-fold copy number amplification of *EGFR* in PC-9 cells increases the size of the mutational target and hence the mutation rate by a factor of eight. In the case of R2, which carries a mutation in NRAS, the copy number of *NRAS* increases the mutation rate by a factor of five. These results apply to a population in the steady state (constant population size) and cannot be compared directly to mutant numbers in recently expanded populations [4], where the number of resistant cells is known to undergo large fluctuations [27, 28].

### The resistance mechanisms are based on different mutations and respond to different compounds

We performed whole exome sequencing and RNA sequencing on all resistant cell lines. R1 exhibits the well-known EGFR T790M mutation [15, 16], and is sensitive to the third-generation EGFR inhibitor osimertinib [29] (see Fig. 2A and SI C for mutation data and detailed response curves).

**Figure 2:**
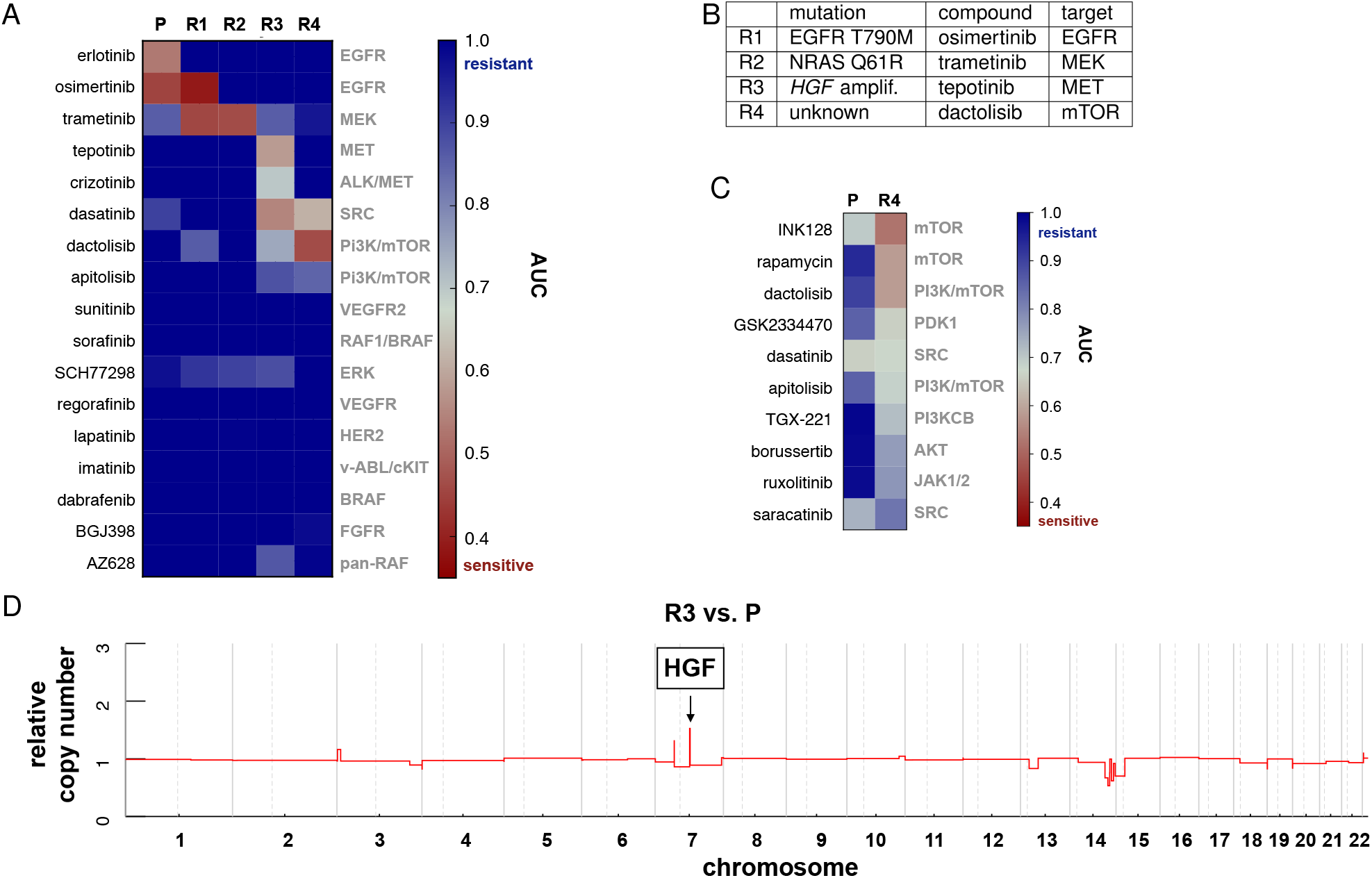
A Drug responses of the parental PC-9 cells and the resistant lines R1-R4 against a panel of compounds targeting potential resistance mechanisms (SI A). High susceptibilities, quantified by the area-under-the-curve (AUC), are indicated in red and compounds on this panel are given in addition to compounds identified in previous iterations, see SI A. Only two compounds, the MEK-inhibitor trametinib and the multikinase inhibitor dasatinib, are effective against more than one resistant cell line. B A summary of the the mutations found in the resistant lines R1-R4, the compounds these lines are susceptible to as identified in panel A, and the targets of these compounds. C Response of R4 to an extended drug panel targeting specifically components of the PI3K/AKT/mTOR pathway. The parental cells are treated with the indicated compounds only, whereas in R4 we measure the additional effect compared to the medium in which R4 was grown, which contains erlotinib, osimertinib, trametinib and tepotinib (EOTT, concentrations 100 nM, 20 nM, 10 nM, 100 nM, respectively). R4 is sensitive to mTOR and PI3K/mTOR dual inhibition, but not to the inhibition of AKT. D R3 shows a focal amplification of the MET ligand *HGF* (length of amplification: 1200 kb).

R2 exhibits a point mutation in NRAS, Q61R, whereas the EGFR T790M mutation is absent. Mutations in NRAS Q61 are known activating oncogenic mutations [30], confer sensitivity to MEK inhibition [31], and can cause resistance to EGFR inhibition in PC-9 cells [32]. Indeed, R2 is most sensitive to the joint inhibition of EGFR and MEK using the MEK inhibitor trametinib (Fig. 2A).

In R3, we found a focal amplification of the growth factor *HGF* (Fig. 2D), and an overexpression of HGF and MET (SI E). Exposure to HGF has been shown to reduce the sensitivity of an EFGR mutant cell line to inhibition of EGFR [33]. HGF is the ligand of MET, a growth factor receptor whose amplification can lead to acquired resistance to EGFR inhibition [26, 1]. R3 is most sensitive to the inhibition of MET (jointly with the inhibition of EGFR and MEK).

In R4, we did not find an obvious genetic alteration responsible for resistance. Chromosome 3 in R4 exhibits a broad (40Mb) amplification up to ten-fold relative to the parental cells (SI C to E). R4 is susceptible to the dual PI3K/mTOR inhibitor dactolisib (jointly with inhibition of EGFR, MET and MEK). To better understand which part of that pathway needs to be inhibited, we screened R4 against an extended panel of compounds (Fig. 2C and SI A). We find that R4 is sensitive to mTOR inhibition and to PI3K/mTOR dual inhibition, but not to inhibition of AKT or SRC. A resistance mechanism based on mTOR signaling has been previously found in EGFR-activated cell lines upon joint inhibition of EGFR and MEK [34], and also in PC-9 cells [9]. Further, the activation of PI3K signaling as an escape mechanism to EGFR inhibition has been observed in different studies [35, 36]. To test if the different cell lines maintain their resistance without selection pressure, we cultivated the four resistant lines without treatment and repeatedly measured their response to erlotinib (SI C). No significant reduction in resistance was found even after 14 weeks of drug withdrawal, implying stable genetic or epigenetic resistance mechanisms.

### The resistance mechanisms show distinct disruptions in signaling pathways

To probe how the different resistance mechanisms affect cellular signaling, we performed immunoblotting of key components of the EGFR signaling pathway in the parental and resistant lines. In the parental line P, treatment with erlotinib shuts down both phosphorylation of EGFR (pEGFR) and also downstream pERK signaling (Fig. 3A). In R1, the delivery of erlotinib does not affect pEGFR or any downstream signaling. This is explained by the T790M mutation, which sterically prevents binding of erlotinib within the ATP binding pocket of EGFR [15, 16]. Osimertinib instead binds covalently to the EGFR receptor [37] and indeed inhibits the phosphorylation of EGFR and the pERK signal. In R2, the pEGFR signal is shut down both by erlotinib and osimertinib, but it does not translate into inhibition of ERK signaling. This is compatible with the activating NRAS mutation Q61R acting downstream of EGFR. The additional treatment with trametinib inhibits MEK and shuts down the pERK signal (Figure 2A and SI A).

**Figure 3:**
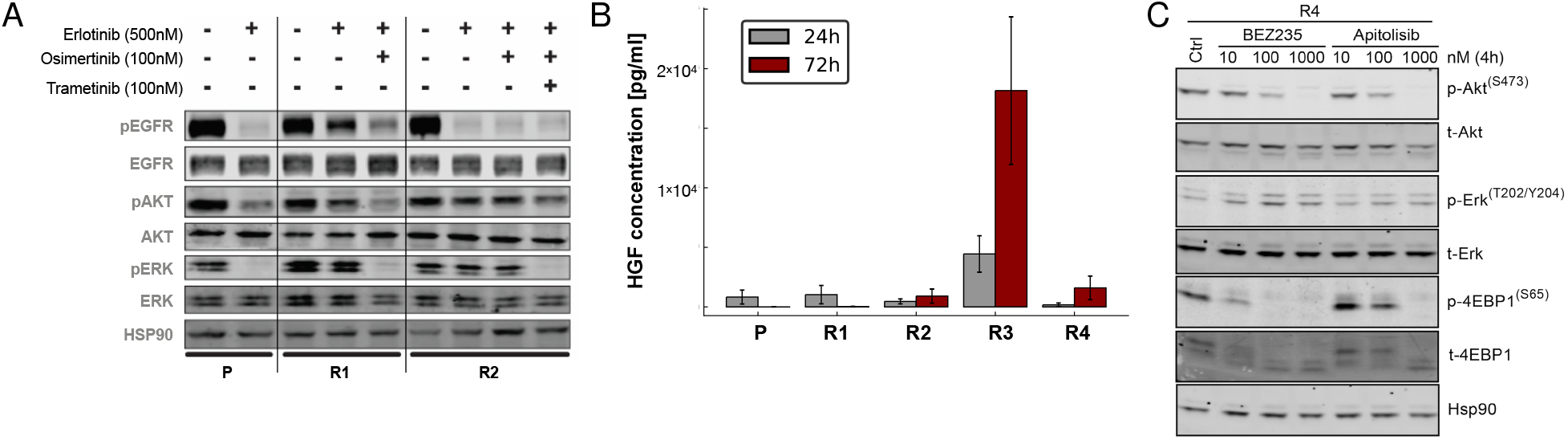
Cell signaling in the resistant cell lines R1–R4. A Protein levels show distinct disruptions of cell signaling in R1 and R2 cells. In R1, the third-generation EGFR inhibitor osimertinib silences EGFR and the downstream ERK signal. In R2, the additional delivery of the MEK inhibitor trametinib is required to silence pERK. B A HGF-specific ELISA shows elevated HGF levels in the R3 supernatant, compatible with the *HGF* amplification found in Fig. 2D and MET/HGF overexpression found by RNA-Seq, see SI E. C Western blots of components of the PI3K/AKT/mTOR pathway in R4 show that neither dactolisib (BEZ235) nor apitolisib, both of which R4 is sensitive to, affect the pERK signal. Instead, they shut down p4EBP1 (a downstream target of mTOR) and pAKT. The housekeeping gene HSP90 serves as a control and the treatment is to be understood as EOTT as in Fig. 2C plus the indicated compound.

In R3, the amplification of *HGF* led to led to an increased concentration of HGF in the supernatant of R3 relative to all other lines, which was assessed using an ELISA assay for HGF (Fig. 3B). It has been previously shown that a high concentration of the growth factor HGF can induce resistance to EGFR inhibition in EGFR-driven cancer cell lines [33]. To validate these observations in PC-9 cells, we treated parental cells with 50 ng/ml of HGF, which lead to an increase of the GI_50_-value for erlotinib of several orders of magnitude (SI E). The finding that R3 is sensitive to MET inhibititors is thus compatible with a MET-bypass track [26] being activated by *HGF* amplification found in R3. In R4, we find that treatment with the dual PI3K/mTOR inhibitors dactolisib or apitolisib does not affect the phosphorylation of ERK, but shuts down both p-4EBP1 (a downstream target of mTOR) and p-AKT (Fig. 3C).

### No further resistance mechanisms arise under a combination of compounds targeting R1-R4

The combination of four compounds erlotinib, osimertinib, trametinib and dasatinib (EOTD) suppresses both the parental cells P, as well as the resistant lines R1-R4 (Fig. 2A). Following our iterative scheme, we attempted to isolate a further resistance mechanism under this drug combination (concentrations 100 nM, 20 nM, 10 nM, and 10 nM, respectively) from populations of size 5 × 10^8^ cells. However, we failed to find any cells growing under EOTD. We then applied the EOTD combination to a population of 5 × 10^7^ cells whose mutation rate had been enhanced approximately hundredfold by a single dose of the mutagen ENU (SI F). Again, no colonies of resistant cells were observed under the EOTD combination. Similarly, no proliferating cells were observed under the combination of erlotinib, osimertinib, trametinib, tepotinib (targeting R3) and dactolisib (targeting R4) (concentrations 100 nM, 20 nM, 10 nM, 100 nM, 100 nM, respectively).

To assess the general toxicity of our EOTD combination we exposed murine IL-3 dependent pro B cells (Ba/F3) and human HEK293T cells to these drugs. We did not observe a significant reduction in cellular viability under EOTD treatment (SI G).

### Drug-tolerant cells survive combination treatment but no longer proliferate

In order to probe if all cells from the original population had been eradicated by EOTD treatment, we withdrew the treatment after seven weeks. Two weeks after the end of treatment, we observed the growth of a small number of colonies. These colonies were in turn eradicated by a dose of 100 nM erlotinib, they thus do not carry a mutation which makes them resistant to erlotinib. This behaviour was reproduced over different treatment periods and we consistently obtained proliferating cells after ending the EOTD treatment (Fig. 4).

**Figure 4:**
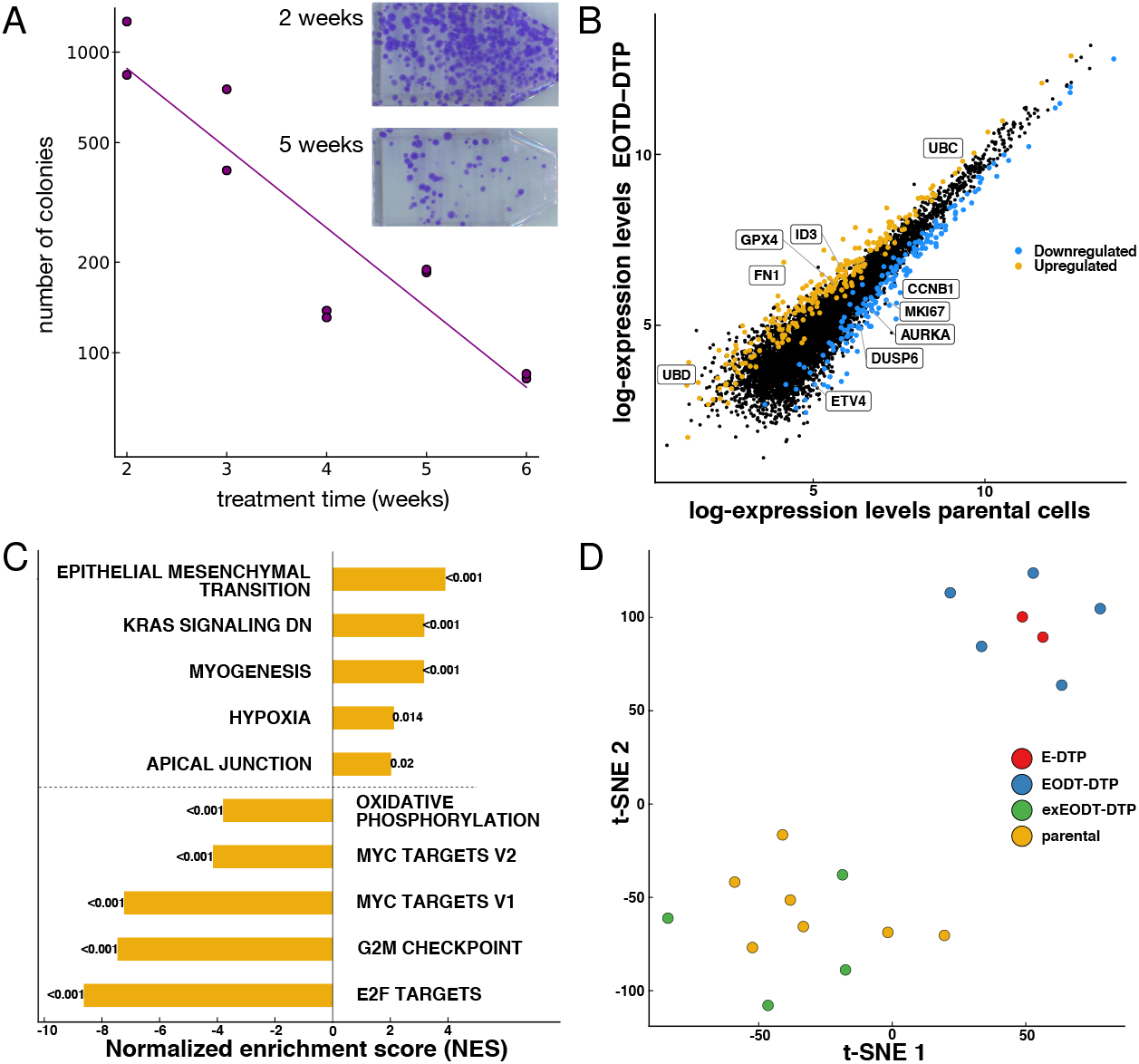
A The number of colonies developed from EOTD-DTP decays exponentially with treatment time. Inset: Colonies developed from EOTD-DTP by first treating flasks containing 5 × 10^7^ cells with EOTD for two weeks (top) and five weeks (bottom), followed in each case by a treatment pause of 2.5 weeks for colony growth and CV staining. The number of colonies is lower for the longer treatment duration as a fraction EOTD-DTP exits the drug-tolerant state during treatment and are thus eradicated. Each data point in the figure stems from such an experiment but with different treatment durations. The line gives an exponential fit with a half-life of 1.13 ± 0.08 week. B Gene expression of drug tolerant cells measured by RNAseq compared to parental PC-9 cells. Differentially regulated genes were identified by an absolute fold change greater than two and a *p*-value < 0.01 (adjusted for multiple testing). C Hallmark gene sets which are upregulated (top) and downregulated (bottom) in EOTD-DTP compared to parental cells. D t-SNE analysis [40] of the expression of the 2000 most variable genes shows that EOTD-DTP (red) cluster with E-DTP (blue). On the other hand, EOTD-DTP proliferating again after the the end of treatment (exEOTD-DTP, green) cluster with parental cells (yellow).

Drug tolerant persisters (DTP) form a subpopulation of cells in a reversible cellular state which allows them to survive under drug treatment without specific resistance mutations [25, 4, 9]. We hypothesize that the behaviour we observe is due to DTP; specifically that some of the DTP exited the drug tolerant state after the treatment ended. These cells grow colonies but are susceptible to erlotinib. Cells exiting the drug tolerant state before the treatment ends, on the other hand, are quickly eliminated. Under this scenario, we expect that the number of DTP decreases with the duration of EOTD treatment.

We distinguish the DTP by the environment they were cultivated in, EOTD or erlotinib. To quantify the dynamics of EOTD-DTP exiting the drug-tolerant state we vary the duration of EOTD treatment and track the number of colonies which emerge after the end of treatment. Upon treating a population of 5 × 10^7^ PC-9 cells with EOTD over different time periods, the surviving cells were kept in regular medium for 2.5 weeks, a time in which a fraction of them exited the drug-tolerant state and formed colonies. These colonies where stained with crystal violet (CV) and counted. The number of colonies decreases with duration of the EOTD treatment and can be fitted to an exponential decay curve, see Fig. 4A. The exponential depletion of the number of EOTD-DTP points to them stochastically exiting the drug-tolerant state at a constant rate. From the decay curve we infer that the drug tolerant state has a half-life of 1.13 ± 0.08 weeks and that the number of EOTD-DTP present at the start of the treatment is approximately 1000 cells in a population of 5 × 10^7^, or 1 in 50000. We checked that the depletion of EOTD-DTP with time was not caused by changes of the medium by repeating the experiment with medium changes at twice the frequency (SI G).

We observe that under EOTD treatment, flasks contain isolated cells at a lower density than under erlotinib, rather than colonies of growing cells, see SI H for microscope images. We conclude that while these cells are drug-tolerant, they do not proliferate under EOTD treatment. They thus behave differently from DTP cultivated under erlotinib (E-DTP), which have been intensively investigated recently [25, 4, 9]. E-DTP proliferate slowly, allowing them to eventually acquire resistance mutations [4]. Conversely, the EOTD-DTP do not proliferate, which curtails their ability to acquire resistance mutations. Consequently, when EOTD-DTP exit the drug-tolerant state, they are sensitive to erlotinib. As a result, these cells die under EOTD treatment, in the absence of treatment they form erlotinib-sensitive colonies. No significant difference in erlotinib sensitivity was found between the colonies that emerged after ending EOTD treatment and the parental PC-9 cells, see SI C.

We ask if there are distinct cellular states underlying the different behaviour of EOTD-DTP and E-DTP. We measured gene expression levels of EOTD-DTP and E-DTP using 3’ UTR RNA-Seq, and compare them to parental PC-9 cells and to EOTD-DTP which have returned to a proliferating state after the end of treatment (exEOTD-DTP), see Fig. 4B–D. A gene-set enrichment analysis comparing EOTD-DTP with parental PC-9 cells shows overexpression in genes associated with the epithelial–mesenchymal transition (EMT), KRAS signaling, and myogenesis, and downregulation in targets of MYC and E2F, as well as genes linked to the G2M checkpoint signature (*p* < 0.001, see Fig. 4C). A signature of high mesenchymal state has been found previously in drug-tolerant persisters derived from different cell lines [38, 39]. To see if gene expression patterns of the different DTP-types fall into separate clusters we use t-distributed stochastic neighbour embedding (t-SNE) [40]. We find that expression levels of exEOTD-DTP cluster with parental cells, but separately from EOTD-DTP (Fig. 4D). On the other hand, expression levels of EOTD-DTP cluster with E-DTP, pointing to the absence of genome-scale differences in gene expression between EOTD-DTP and E-DTP.

To further probe the distinction between EOTD-DTP and E-DTP, we switched individual DTP populations between erlotinib treatment and EOTD treatment. We find that EOTD-DTP start proliferating once the treatment is switched from EOTD to erlotinib (Fig. 5A). Conversely, E-DTP growing under erlotinib are rapidly depleted when switching to EOTD (Fig. 5B). We conclude that drug tolerant persisters proliferate under erlotinib [4, 9], they do not proliferate under EOTD, irrespective of how they were generated. Taking these observations together, we conclude that despite their different behaviour, EOTD-DTP and E-DTP are not distinct entities. Instead, the different behaviour under erlotinib and EOTD is due to the different drug regimes.

**Figure 5:**
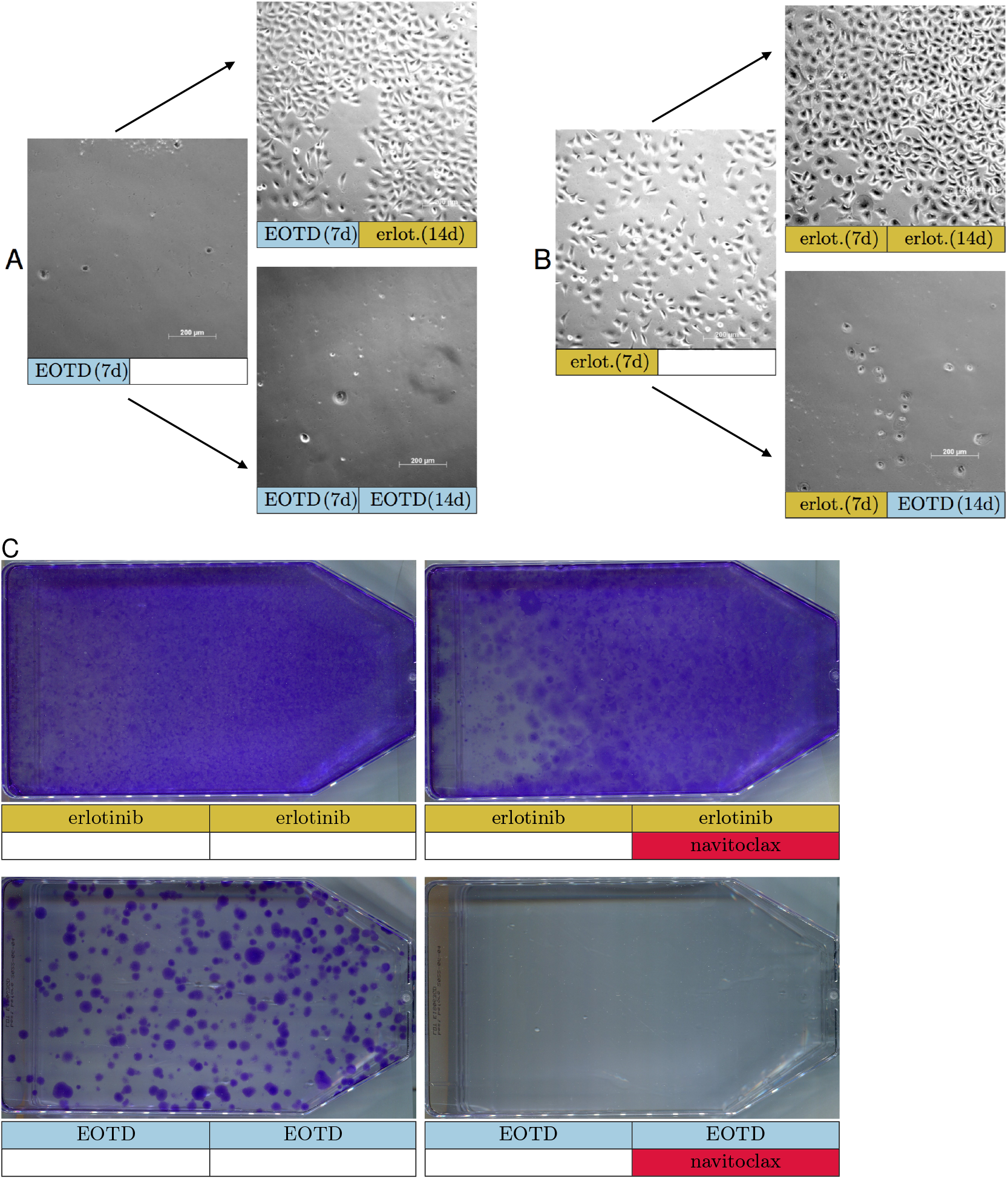
Switching drug-tolerant persisters between erlotinib and EOTD treatment. A We plated 7 × 10^5^ cells kept in regular medium for one day and then treated with EOTD for one week (left). This treatment was then switched to erlotinib for two weeks (top right) or maintained on EOTD as a control (bottom right). B Analogously, but initial treatment was with erlotinib for one week (left) and switched to EOTD for two weeks (bottom right) or maintained (top right). Images are representative of 4 independent experiments. We find that DTP proliferate under erlotinib, but not under EOTD. We checked that populations growing under erlotinib did not contain the mutation EGFR T790M. C Eradicating a large population of PC-9 cells with EOTD and navitoclax: PC-9 cells were plated in T175 flasks and left untreated until confluency. The flasks were then treated for two weeks with erlotinib (top) or with EOTD (bottom). In the flasks on the right, 1 *μ*M navitoclax was added during the second week. All flasks were CV stained following a drug holiday of 2 weeks.

### Targeting both resistant and drug-tolerant cells

Finally, we ask how a large population of PC-9 cells responds to suppressing R1–R4 using the EOTD combination and also eradicating DTP. The latter have previously been shown to be sensitive to the BH3-mimetic Bcl-2-inhibitor navitoclax [4]. We cultivate populations of approximately 5 × 10^7^ PC-9 cells (to allow for visual and microscopic inspection of the population under therapy) and treat two arms with erlotinib and two arms with EOTD for two weeks, either with and without 1 *μ*M navitoclax in the second week. After the end of the treatment, we waited for two more weeks before staining. Under erlotinib, a dense layer of cells is found (Fig. 5C), presumably due to both resistant cells which repopulate the flask after the end of treatment, as well as cells having gone through the drug-tolerant state. Under erlotinib and navitoclax, this layer is formed by from erlotinib-resistant cells only. Under EOTD without navitoclax, hundreds of individual colonies are seen, similar to Fig 4A. We interpret these as DTP which returned to a proliferating state after the end of treatment. Only under EOTD *and* navitoclax not a single colony is seen. This behaviour was consistent over three replicates. We conclude that EOTD and navitoclax can eradicate a large population of PC-9 cells entirely. To assess the toxicity of the EOTDN combination, we measured the viability of cell lines not driven by an EGFR mutation under EOTDN. We find no significant viability reduction due to the EOTDN combination in the human cell line HEK293 and the murine Ba/F3 show, see SI G.

## 3 Discussion

The rapid emergence of resistance to targeted therapy in solid tumors is owed to the combination of a large number of proliferating cells and a finite mutation rate. Together, these generate a large supply of resistant mutants, which expand under therapy. Resistance is thus avoided under only a limited number of well prescribed scenarios. The first is a small population size of proliferating tumor cells, such that the tumor is unlikely to contain a resistant cell at the start of therapy. However, solid tumors typically consist of many millions of cells at discovery, and thus generally contain resistant cells [1, 2, 3, 4]. The early detection of smaller tumors, for instance through routine screening of DNA circulating in the blood [41], may change this in the future. The second scenario is a cell requiring several mutations to become resistant. If a (hypothetical) tumor depended on two oncogenic mechanisms, a dual therapy would require two independent resistance mutations to cause acquired resistance to both therapies [42, 43]. Such double mutants do typically not appear as part of the standing genetic heterogeneity at realistic mutation rates and population sizes [43]. While no tumors susceptible to such a dual targeted therapy have been found so far, cancer immunotherapy may fall into this conceptual category, with the immune system responding to multiple neoantigens.

Our work is a proof-of-principle study for a third approach: treating all pre-existing resistance mechanisms as part of first-line therapy, either simultaneously or in quick succession. We have shown that in an EGFR-mutant cell line, PC-9, four compounds are sufficient to suppress all resistance mechanisms existing in a population of 5 × 10^8^ cells. This is surprising given the large number of potential resistance mechanisms to EGFR inhibition [44]. The resistance mechanisms which need to be suppressed comprise a secondary mutation in EGFR (the well-known gate-keeper mutation T790M), the activation of signaling downstream of EGFR by an NRAS mutation, and the activation of alternative bypass pathways. These mechanisms all coexist in our population at the onset of therapy, and will proliferate under any therapy which does not inhibit them. Also, we have found that the different resistance mechanisms grow at different rates, with EGFR T790M growing fastest, and NRAS-mutant and *HGF*-amplified mechanisms growing more slowly. The growth rate of a resistance mechanism, along with mutation rate, determines the population fraction of cells with a given mechanism. As a result, it may be that cells with other mechanisms grow too slowly to be present in our population, but would appear in the standing genetic heterogeneity of an even larger population.

The growth rates of the different resistance mechanisms are likely to depend on their genetic background. Depending on how much these growth rates differ across cell lines and across tumors, different cancer cell populations might contain different resistance mechanisms, even if caused by the same oncogenic driver. Or, alternatively, populations with the same driver mechanism might contain largely the same set of resistance mechanisms. Differences in growth rates will then still affect the response to therapy. Specifically, the mechanism with the highest growth rate might differ across populations. This would lead to different mechanisms emerging under therapy, even if the resistance mechanisms present in the population were largely the same. The extent to which the spectrum of resistance mechanisms is shared across different tumors with the same oncogenic driver is an important question for further research. Our simple iterative scheme to amplify rare resistant mutants can answer this question in the context of cell cultures and ultimately help inform treatment options.

The growth rates of different resistance mechanisms differ by at most 20% from one another. The benefit of eliminating one or even several resistance mechanisms, while leaving others untreated, is thus small: The time it takes for another, untreated mechanism to emerge is comparable to the time it would have taken for one of the treated mechanisms to emerge. This provides a rationale for the current treatment paradigm, where a first-line treatment is continued until the emergence of a resistant tumor, and is then switched; the time gained by waiting several times for the emergence of different resistant mutants exceeds the time gained from waiting once for the emergence of a slightly slower-growing mechanism. It may also account for the failure of drug combinations targeting a subset of resistance mechanisms to increase progression-free survival by more than a few months in clinical trials on solid tumors [45, 46, 47, 48].

Inhibiting all resistance mechanisms present in a population, as well as the parental cancer cells leaves a population devoid of rapidly growing cells. However, such a population can still contain drug-tolerant persisters. DTP have been studied extensively in PC-9 cells [25, 4, 9, 39]; they form a subpopulation of cells growing slowly under erlotinib, and can eventually acquire resistance mutations. Interestingly, the EOTD combination, which inhibits the different resistance mechanisms in PC-9 cells, prevents the expansion of drug-tolerant persisters as well. Instead of slowly expanding, the number of DTP exponentially decays under EOTD with a half-life of approximately one week. This lack of growth severely limits the potential of DTP to acquire resistance mutations. Also, the DTP can be eliminated using BCL2 inhibitors such as navitoclax [4, 9]: combining erlotinib, osimertinib, trametinib, dasatinib (EOTD) with navitoclax cleared flasks of ca. 5 × 10^7^ cells, whereas erlotinib alone or with navitoclax, or EOTD alone lead to colony formation after the treatment ended (Fig. 5C).

The result that different resistance mechanisms require different drugs specific to the particular mechanism points to an important distinction between resistance to targeted cancer therapy and resistance to antibiotics or to antiviral drugs. For example, in HIV, resistance to a single anti-retroviral drug emerges quickly, again due to the high viral mutation rate and population size. However, cocktails of drugs targeting different aspects of the virus have been used successfully for decades, leading to a nearly normal life-expectancy of HIV patients [49]. Similarly, antibiotics targeting different surface proteins, or different aspects of the bacterial metabolism have been combined to combat the emergence of resistance to a single antibiotic. This has enabled to cure tuberculosis despite the rapid emergence of resistance to monotherapy [50]. In these cases, viruses or bacteria resistant to one particular drug are sensitive to some (any) other drug of the combination.

The situation in targeted cancer therapy is different: Fig. 2 shows that the different resistance mechanisms require drugs specific to each mechanism. This difference may arise because there is a vast number of independent targets antiviral or antibiotic therapies can exploit, but cancer cells typically do not have multiple independent oncogenic dependencies. As a result, each resistance mechanism present at the start of therapy requires a specific compound. In the corresponding case in antibiotics/antivirals, if a mutation confers resistance to attacks on one particular target, a different molecular target of the pathogen can be chosen (rather than seeking to diable the particular resistance mechanism).

In this context, it is encouraging that the spectrum of resistance mechanisms found in a large population of PC-9 cells can be eliminated with a cocktail of five clinically-approved compounds. Future work will exploit that the different resistance mutations arise in different cells, raising the possibility of dosing schedules targeting the different resistance mechanisms in turn while minimizing cross-toxicity.

## Supporting information

Single file Supplementary Information

## Acknowledgment

This work was supported by the German Federal Ministry of Education and Research (BMBF) within the framework of the e:Med research and funding concept (grant e:Med SMOOSE), by the DFG SFB 1310, by the German federal state North Rhine-Westphalia (NRW) as part of the EFRE initiative (grant LS-1-1-030a to M.L.S.), the Deutsche Krebshilfe (70112888 to M.L.S. and 70113129 to C.L.) and the Else Kröner-Fresenius Stiftung (Memorial Grant 2018_EKMS.35 to J.Br.). Borussertib [51] and other compounds were kindly provided by the Rauh lab, University of Dortmund. Ba/F3 cells were a kind gift from Nikolas von Bubnoff.

## 4 Materials and Methods

### 4.1 Cell culture

PC-9 cells were cultured in RPMI 1640 (ThermoFisher, USA) supplemented with 10 % FBS and 1 % penicilin-streptomycin. R1 cells were cultured in the presence of 500 nM erlotinib; R2 in the presence of 500 nM erlotinib, and 100 nM osimertinib; R3 in the presence of 100 nM erlotinib, 20 nM osimertinib, and 10 nM trametinib; R4 in the presence of 100 nM erlotinib, 20 nM osimertinib, 10 nM trametinib, and 100 nM tepotinib. The identity of the parental and all resistant cell lines was confirmed by STR finger printing. All cell lines were checked for mycoplasma.

### 4.2 Growth rate measurements

20000 cells per well were plated in 12 well plates. After approximately 24 h, 48 h, 72 h, and 96 h the number of cells per well was counted (three wells per time point). The growth rate *g* was calculated by fitting the resulting numbers to an exponential growth curve *N* (*t*) = *N*_0_ exp(*gt*). The experiment was performed at least three times per cell line. The reported growth rates are given as the mean plus/minus the standard error over independent runs of the experiment.

### 4.3 Cell viability measurements

1500 cells per well were plated in 96 well plates in triplicate for each condition. After 24 h the wells were treated with the respective drug or drug combination. After 96 h of expansion cell viability was measured using CellTiter-Glow assay (Promega, USA) following the manufacturer’s instructions and normalized with respect to a control grown under DMSO. All measurements were repeated in triplicate.

For clonogenic survival assays of DTP, cells were seeded and left to adhere over night before start of treatment. Cells were treated with erlotinib or EOTD for one week followed by one week of combinatorial treatment with erlotinib or EOTD with 1*μ*M of indicated compounds. After treatment, compound containing medium was removed and replaced by culture medium to allow for viable cells to grow out. Adherent cells were fixed using 4% paraformaldehyde and stained using 0.1% crystal violet solution.

### 4.4 DNA-seq

We extracted cellular DNA using the Gentra Puregene Tissue Kit (Qiagen, Germany) following the manufacturer’s instructions.

Whole-exome sequencing was performed from genomic DNA using the SureSelect Human All Exon V6 (Agilent, Santa Clara, CA, USA) according to the manufacturerâĂŹs instructions. Obtained exome libraries were paired-end sequenced a HiSeq4000 (2 × 75 bp) platform (Illumina, San Diego, CA, USA).

To confirm the EGFR T790M mutation in R1 and the NRAS Q61R mutation in R2 we performed Sanger sequencing on *EGFR* exon 20 (forward primer TGA AAC TCA AGA TCG CAT TCA, reverse primer ACA CAT ATC CCC ATG GCA AA) and *NRAS* exon 3 (forward primer ACA CCC CCA GGA TTC TTA CAG, reverse primer CAC AAA GAT CAT CCT TTC AGA GAA).

### 4.5 Immunoblotting

For Western Blot analysis cells were seeded and left to adhere over night before treatment with denoted compounds. Cells were harvested after 4 hours of treatment and lysed in RIPA buffer (150 mM NaCl, 50 mM Tris/HCl [pH 7.5], 1% Triton-X 100, 0.5% Na-Deoxycholate, 0.1% SDS) supplemented with protease inhibitors (Roche complete mini) and benzonase (Millipore). Proteins were separated on 4 − 12% Novex Tris-glycine gels (Invitrogen), and transferred to PVDF membrane (Millipore). Membranes were blocked using TBS with 2% Cold Water Fish Gelatine (Sigma-Aldrich) and incubated with primary antibodies in blocking solution with 0.2% Tween 20 overnight at 4 °C. Membranes were washed in TBS-T, incubated with secondary antibodies for 1 hour at room temperature and washed in TBS-T before detection with Odyssey CLx imaging system (LI-COR).

Primary antibodies: p-EGFR (CST #3777, 1:1000), EGFR (CST #2232, 1:1000), p-Erk (CST #4370, 1:1000), Erk (CST #9102, 1:1000) p-Akt (CST #4060, 1:1000), Akt (CST #C2927, 1:1000), p-MET (CST #3077, 1:1000), MET (CST #3148, 1:1000), HSP90 (CST #4877, 1:1000). Secondary antibodies: goat anti-rabbit 800CW (Li-Cor Cat. 926-32211, 1:10000), goat anti-mouse 800CW (Li-Cor Cat. 926-3220, 1:1000

Hepatocyte Growth Factor (HGF) concentrations in the supernatant were measured at indicated time points after seeding the cells in 12-well plates. Measurements were performed using the Quantikine ® human HGF immunoassay (R&D Systems) according to the manufacturer’s instructions.

### 4.6 RNA-seq

For PC-9 parental and R1-4 derivatives RNA was isolated using the RNeasy Mini Prep Kit (Qiagen, Germany). Due to the low number of EOTD-DTPs, RNA extraction from EOTD-DTP was performed with the Arcturus PicoPure RNA-isolation kit (Thermo Fisher, USA). For details see SI E.

### 4.7 Colony formation of exEOTD-DTP

10^6^ cells per flask were plated in T175 culture flasks in normal medium. After one week, treatment with EOTD was started and maintained for the indicated time periods. Treatment was withdrawn and cells were cultured in normal medium for 2.5 weeks. Emerging colonies were stained with crystal violet and counted. The resulting values were fit to an exponential curve. The medium was changed twice per week throughout.

