## Supplementary material for "In vitro model for resistance in oncogene-dependent tumors at the limit of radiological detectability": Single file Supplementary Information

September 3, 2019

#### A The drug panel and dose-response curves

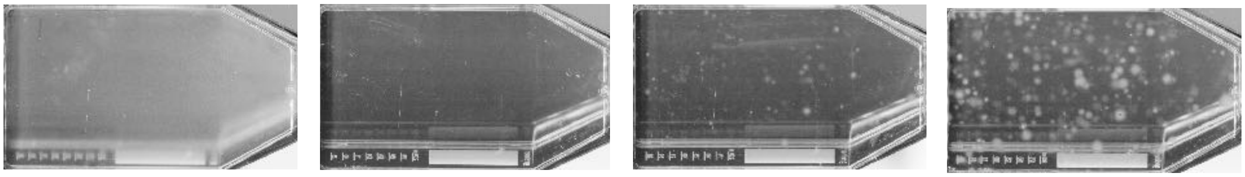

**Figure 1:** Visualizing the emergence of resistant cells in a large population under targeted treatment. For illustration purposes, PC9 cells are grown in a T175 flask (ca. $5 \times 10^7$ cells). Upon confluence they were exposed to 500 nM of erlotinib. Pictures show the flask on days 1, 4, 14 and 18 of treatment (left to right). At the start of treatment, the whole growth area is filled with cells, leading to an opaque appearance. Already on day 4 the flask appears clear, however 14 days into treatment, small colonies become visible and start to expand despite the ongoing exposure to erlotinib.

Each resistant lines was grown starting from a population of ca. $5 \times 10^8$ PC-9 cells in a hyperflask (Corning, USA) with a growth area of $10 \times 175\,\mathrm{cm}^2$ under constant drug concentration. Surviving cells are trypsinated and transferred to a T175 flask after approx. 2 months. The constant drug regime was maintained until 80% confluence was reached, when cells were trypsinated again and taken for analysis. Each iteration took $12 - 15$ weeks depending on the growth rate of the resistant line.

We screened the resistant cell lines against the following drug panel:

| **compound** | **target** | **compound** | **target** |
|---|---|---|---|
| Erlotinib | EGFR | AZ628 [1] | pan-RAF |
| Osimertinib | EGFR (T790M) | Imatinib | v-Abl/c-Kit/PDGFR |
| Lapatinib | EGFR/HER2 | Sorafenib | RAF-1/BRAF/VEGFR-2 |
| Tepotinib | MET | SCH77298 [2] | ERK1/2 |
| Crizotinib | MET/ALK | Sunitinib | VEGFR2/PDGFR$\beta$ |
| Trametinib | MEK | Regorafinib | VEGFR1/VEGFR2/VEGFR3/PDGFR$\beta$/Kit/RET/RAF1 |
| Dabrafenib | BRAF (V600) | Dactolisib | PI3K/mTOR |
| Apitolisib | PI3K/mTOR | Dasatinib | Abl/SRC/c-Kit |
| BGJ398 [3] | FGFR | | |

The resistant cell line R1 was derived under 500 nM erlotinib. R2 was derived under 500 nM erlotinib and 100 nM osimertinib. To decrease cross-toxicity we reduced the concentration for the other resistant cell lines: R3 was derived under 100 nM erlotinib, 20 nM osimertinib and 10 nM trametinib. R4 was derived under 100 nM erlotinib, 20 nM osimertinib, 10 nM trametinib and 100 nM tepotinib. These concentrations were chosen to provide at least five times the IC-50 levels of the preceding line, e.g. the IC50 for parental cells under erlotinib is approx. 100 nM, so R1 was grown under 500 nM erlotinib to rapidly deplete the population of all cells not carrying a resistance mechanism against erlotinib, and analogously for the subsequent iterations.

We measured cell viability $v(c)$ at three different concentrations $(10, 100, 1000\,\text{nM})$. To quantify the sensitivity of a cell line to a particular drug, we calculated the area under the curve as

$$\text{AUC} = \frac{v(10\,\text{nM}) + v(100\,\text{nM}) + v(1000\,\text{nM})}{3}. \tag{1}$$

R1 was screened against all elements of the drug panel in medium only; R2 was screened under treatment of 500 nM erlotinib; R3 under 100 nM erlotinib, and 20 nM osimertinib; and R4 under 100 nM of erlotinib, 20 nM osimertinib, and 10 nM trametinib. For each cell line, we chose the drug with the lowest AUC as the most effective drug. Only in the case of R3 we used tepotinib instead of dasatinib to derive further resistant lines even though the latter had a lower AUC: no further cells grew under erlotinib, ostimertinib, trametinib, dasatinib (EOTD, see SI G). EOTD concentrations were 100 nm erlotinib, 20 nM osimertinib, 10 nM trametinib and 10 nM dasatinib throughout. To confirm the result of the initial screening we tested the drugs that showed highest sensitivity at a higher resolution (with concentrations ranging from 1 nM to 10 $\mu$M). The detailed response curves for the compounds R1-R4 respond most strongly to are given in Fig.2.

The extended drug panel used in Fig 2C was chosen to probe different components of the AKT-mTOR pathway. INK128 is sapanisertib targeting mTOR [4], GSK2334470 targets PDK1 and is characterized in [5], and TGX-221 targets PI3KCB [6]. Borussertib targets AKT [7] and was kindly provided by the Rauh lab, University of Dortmund.

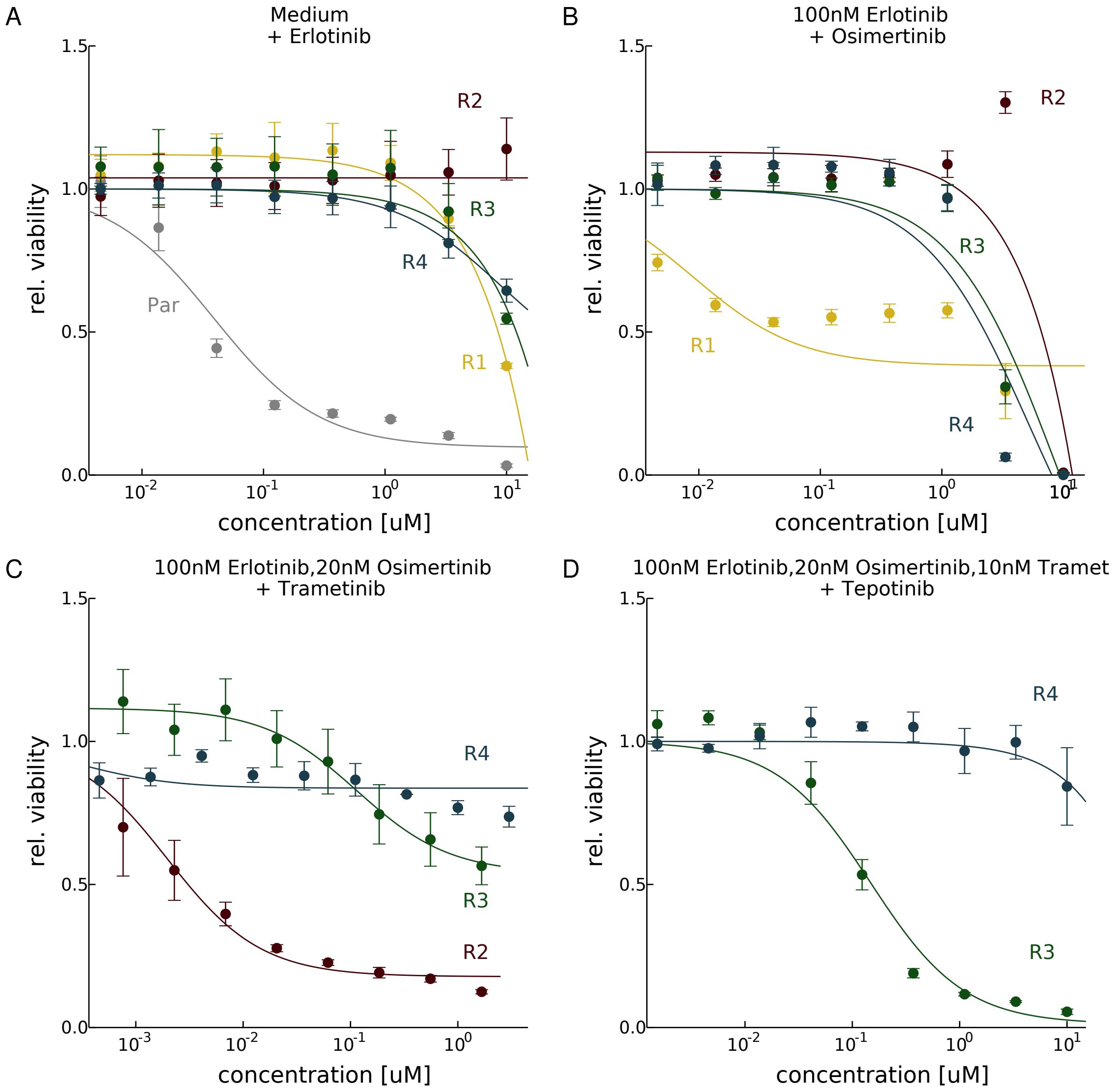


**Figure 2:** Detailed dose-response curves for the compounds the cell lines respond most strongly to. The parental cell line is indicated in gray, R1 in yellow, R2 in red, R3 in green, and R4 in blue.

#### B Population genetic model and parameter estimates

We consider a population of cells with a finite carrying capacity. The carrying capacity $K$ limits the total number of cells in the population, for instance due to a finite supply of nutrients or due to spatial constraints. This differs from models of exponentially growing populations considered previously [8].

There are two types of cells, cells which are sensitive and and cells which are resistant to a particular targeted therapy. We are interested in the statistics of the number of resistant cells given the size of the population, the mutation rate from sensitive to resistant cells, and growth and death rates. We focus on the case where the majority of the cells is sensitive. As their number is large, we model their dynamics by a deterministic growth law. Against this background, we consider the stochastic emergence and growth of resistant mutants.

The process starts with $N_0$ sensitive cells which divide at rate $b_s(t) = b_s\left(1 - \frac{N_s(t)}{K}\right)$, die at rate $d_s$, and mutate at rate $\mu$ per cell division to a resistant cell. Solving this deterministic growth model yields

$$N_s(t) = \frac{N}{1 + \omega \exp(-g_s t)}, \tag{2}$$

where $\omega = \frac{N}{N_0} - 1$ and $N = K\left(1 - \frac{d_s}{b_s(1-\mu)}\right)$. $N_s(t)$ denotes the number of sensitive cells at time $t$.

Resistant cells emerge stochastically at rate $\mu\, b_s(t)\, N_s(t)$, where $\mu$ is the mutation rate per cell division. Once introduced to the population, resistant cells divide at rate $b_r(t) = b_r\left(1 - \frac{N_s(t)}{K}\right)$ and die at rate $d_r$. For reasonable choices of parameters ($b_s - d_s > b_r - d_r$) and long times, the system reaches a steady state, where the statistics of the number of resistant cells no longer changes with time. In this limit, the number of resistant cells is distributed according to

$$p(k) = \left(1 - \frac{\beta}{\delta}\right)^{\frac{\mu N}{\beta}} \left(\frac{\beta}{\delta}\right)^{k} \frac{\Gamma\left(\frac{\mu N}{\beta} + k\right)}{\Gamma\left(k+1\right)\Gamma\left(\frac{\mu N}{\beta}\right)}, \tag{3}$$

where $\beta = \frac{b_r}{b_s}$ and $\delta = \frac{d_r}{d_s}$ denote the relative birth and death rates, respectively. For the expected number of resistant cells we obtain from (3)

$$R = \frac{\mu}{1-\mu} N (\delta - \beta)^{-1} \tag{4}$$

and for the probability that there is at least one resistant cell in the population

$$p_{\text{res}} = 1 - \left(1 - \frac{\beta}{\delta}\right)^{\frac{\mu N}{(1-\mu)\beta}}. \tag{5}$$

The probability is of crucial interest as it gives the probability that a resistant tumour can emerge from a resistant cell which pre-existed at the time of treatment. All the results given above can be derived from the probability generating function $G(x)$, which for our model is given by

$$G(x) = \left(\frac{1 - \frac{\beta}{\delta}}{1 - x \cdot \frac{\beta}{\delta}}\right)^{\frac{\mu N}{\beta}}. \tag{6}$$

We also study the dynamics of individual resistant colonies (or clones) for the process defined above, where a colony is a set of cells that are introduced to the population by the same mutational event. Again, we treat the number of resistant colonies as a stochastic variable. We assume colonies to be independent of each other: when the number of resistant cells is small compared to the number of sensitive cells, the growth dynamics of one colony does not affect the dynamics of another. Colonies are introduced at the same rate at which new resistance mutation occur $\nu(t) = \mu\, b_s(t)\, N(t)$ and die at a rate $d_c$, which we need to determine. Colony death is a barrier crossing problem, where death of a colony occurs when the number of cells comprising the colony reaches zero. Again, we focus on the steady state. The number of colonies then follows a Poisson distribution as both creation and death of colonies are independent events occurring at constant rates. The expected number of colonies is then given by

$$C = \frac{\nu(t \to \infty)}{d_c}. \tag{7}$$

and the probability of at least a single colony existing follows as

$$p^c_{\text{res}} = 1 - \exp\left(\frac{\nu(t \to \infty)}{d_c}\right), \tag{8}$$

which has to equal (5). From this relationship, we derive the steady state value for the colony death rate as

$$d_c = -d_s \beta \log\left(1 - \frac{\beta}{\delta}\right)^{-1} \tag{9}$$

giving

$$C = -\frac{\mu}{1-\mu}\frac{N}{\beta}\log\left(1 - \frac{\beta}{\delta}\right). \tag{10}$$

The heatmap in Figure 1D shows equation (4) as a function of $\beta$ and $\mu$. We assume $\delta = 1$ for all resistance mechanisms, i.e., the death rates of resistant cells equal the death rate of sensitive cells. This is compatible with the measurements for PC9 wild-type and PC9 EGFR T790M positive cells reported in [8] ($d_{\text{wt}} = d_{\text{T790M}} \approx 0.06\,\text{d}^{-1}$). We thus use

$$\beta_x = \frac{g_x + 0.06\,\text{d}^{-1}}{g_P + 0.06\,\text{d}^{-1}} \tag{11}$$

with $x = \{\text{R1,R2,R3,R4}\}$ to calculate the relative birth rates. We further set $N = 5 \times 10^8$ as total population size. For the mutation rate we use an order-of-magnitude estimate for the somatic point mutation rate of $1 \times 10^{-9}$ per generation and nucleotide [9]. In R1 the mutational target is approximately eight times larger than the background rate due to an amplification in *EGFR*. In R2 we found five copies of *NRAS*, which is why the mutational target is five times larger than the background mutation rate. In R3 resistance is caused by an focal copy number amplification, see Fig. 6 in SI D. As no reliable estimates exist at which rate amplifications occur in the genome we do not fix a value on the $y$-axis. In R4 we do not know if resistance is driven by a point mutation, a gene amplification or another modification. For this reason, again, we also do not assign a mutation rate estimate to this resistance mechanism.

We also calculate the minimal growth rate at which resistant cells would be present on average with a single cell. Resistance mechanisms with a substantially slower growth rate would typically be absent from a population, those with a substantially faster growth rate would typically be present in multiple cells. To determine the minimal growth rate, we set the average number of cells in equation (4) to one and solve for $\beta$. Choosing the same parameters as above in (11) ($\delta = 1$, $N = 5 \times 10^8$ and $\mu = 10^{-9}$), we find $g_{\min} = 0.38\,\text{d}^{-1}$ as the minimal growth rate, about half the growth rate of the parental cells.

#### C Mutations in resistant cell lines and the stability of resistance under drug holidays

We aligned raw sequencing reads to the human reference genome (NCBI build 37/hg19) using the BWA mem aligner (version 0.7.13-r1126). Concordant read pairs were masked as possible PCR duplicates and areas of overlapping read pairs were excluded from analysis in one read. Point mutations were determined by our in-house cancer genome analysis pipeline [10, 11, 12, 13, 14, 15].

The point mutations EGFR T790M in R1 and NRAS Q61R in R2 were also confirmed by Sanger sequencing, see Fig. 3.

R3 harbours a focal amplification on chromosome 7 discussed in SI D. In R4 we did not find an obvious alteration responsible for the resistance to erlotinib, osimertinib, trametinib. R4 shows point mutations in *CROCC,LRRC7,SPRR1A,ARHGAP30,GPR161, PAPPA2, FMN2, USP34, LIMS2, LRP2, CNPPD1, RAB28, IRX2, BTN2A1, TJAP1, MYO6, GPR141, C7orf25, FSCN3, TRIM24, FGL1, COL22A1, FAM73B, SLIT1, HPSE2, LRRC4C, KCNA5, EPS8, ABCC9, KRT75, GIT2, CENPJ, MOK, ATP10A, VPS18, ZNF592, ALPK3, SV2B, EXOSC6, KRT16, NETO1, TLE6, CILP2, UQCRFS1, CEACAM1, PPP1R13L, SIGLEC8, VN1R2, ZNF264, SLC23A2, NCOA6, MYH7B, PLTP, YTHDF1, UCKL1, C21orf7, FRMPD4, NAP1L2, SUCLG1*, and *TFAP2A* copy number alterations in R4 are discussed in SI D and E.

To test if the observed resistance mechanisms are rapidly reversible or whether they are stable, we cultured R1-R4 in regular medium without treatment for several months. Cells were split twice per week and screened against EGFR inhibition at least once every two weeks. As seen in Fig. 4 A, the sensitivity to EGFR inhibition does not drop rapidly, but decreases slowly. The slow decrease is to be expected as mutations can occur which fitter but less resistant genotypes into the population, which subsequently increase in frequency. Nevertheless, even after 14 weeks of drug holiday, the $GI_{50}$ values of all resistant lines are at least 10 times higher than the respective GI50 values of the parental cells, see Fig. 4 B. In contrast to that, cells that emerged from DTPs that survived the EOTD combination show the same sensitivity to erlotinib as the parental line, see the grey triangles in Fig. 4 A. This means that all resistant lines exhibit stable resistance, whereas the DTP phenotype is reversible.

To test the samples in Fig. 5 A and B (drug tolerant persisters under different treatments) for the EGFR T790M mutations, Digital PCR was performed using the QX200 System (Bio-Rad, Hercules, CA). Probes detecting the EGFR T790M mutation (CP2000019) and other consumables were purchased from Bio-Rad (Bio-Rad, Hercules, CA) and used according to manufacturer's instructions with the exception of the annealing temperature, which was increased to 57C. Data Analysis was performed using Quantasoft 1.7.4.

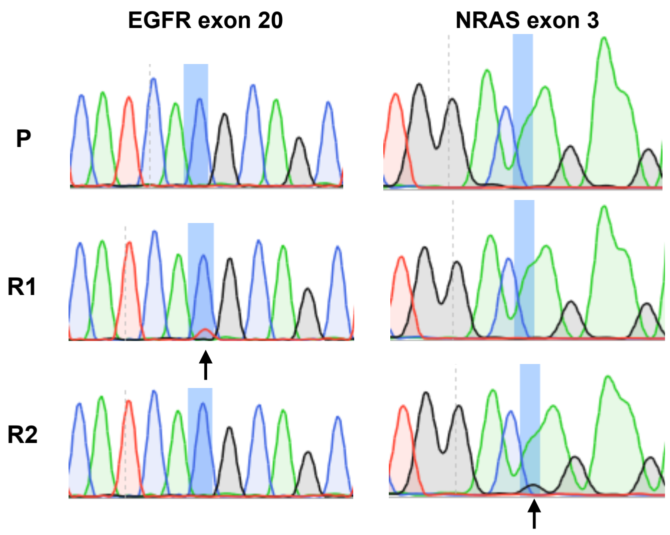


**Figure 3:** R1 exhibits the mutation EGFR T790M and R2 exhibits NRAS Q61R as validated by Sanger sequencing.

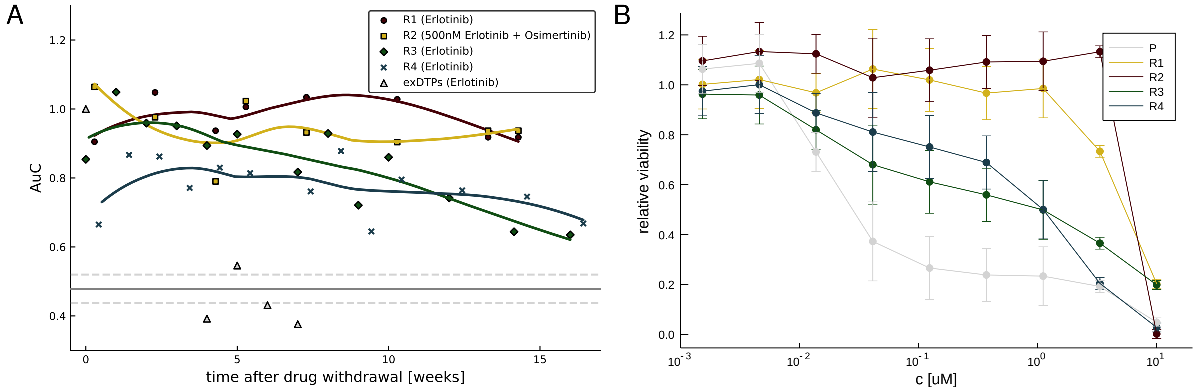


**Figure 4:** A Stability of the resistant lines R1-R4 after compound withdrawal. The resistant lines are tested at different times for sensitivity to EGFR inhibition and the AUC is calculated. The lines indicate the general trend in the data. B Drug response curves for the resistant and the parental cell lines against EGFR inhibition after 100 days of drug withdrawal. R1 and R2 are still fully resistant, whereas R3 and R4 partially lost their resistance (see Fig. 2 for comparison). Nevertheless, the GI50 values remain at least 10 times higher than those of parental cells.

### D DNA copy numbers of parental and resistant lines

To determine the copy number status of from whole exome data (SI C) we used a modified version of Sclust [16] to deal with the absence of a matched normal. The ploidy of PC-9 cells is three.

Fig. 6 shows the copy numbers of the resistant lines R1-R4 versus the parental line. Fig. 7 zooms in on the focal amplification found in R3 containing the *HGF* gene.

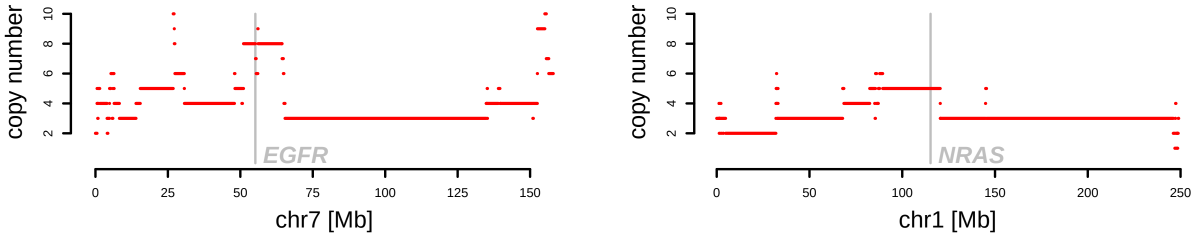


**Figure 5:** Copy numbers of of chromosomes 7 (left) and 1 (right) for the parental PC-9 line. These two chromosomes contain the genes encoding EGFR and NRAS where indicated, respectively. They show an 8-fold broad amplification of the region containing *EGFR* and a 5-fold amplification of the region containing *NRAS*.

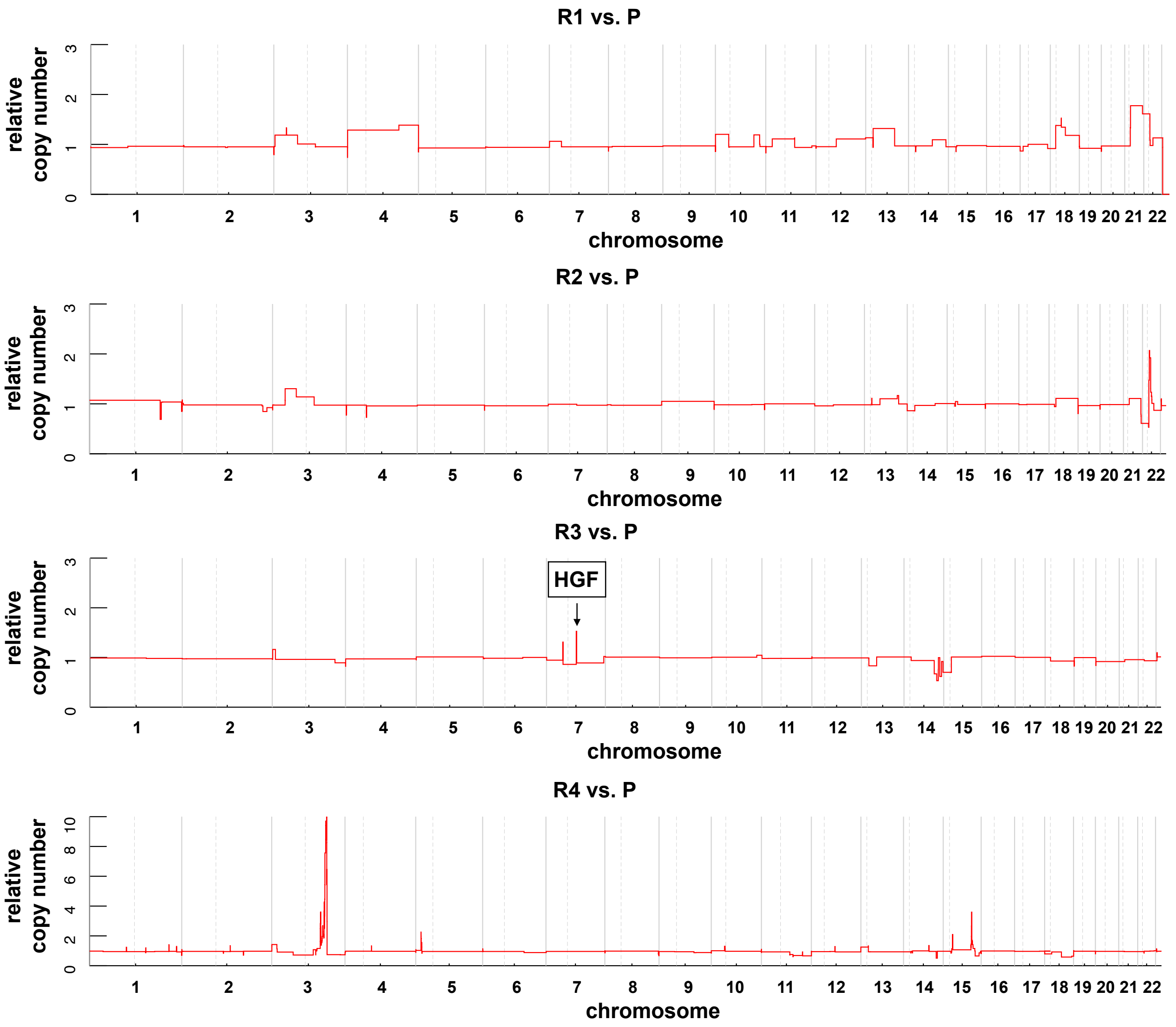


**Figure 6:** Relative copy number of the resistant lines R1–R4 compared to the parental line. Note the scale of the y-axis in R4: there is a broad amplification (40Mb) with relative copy numbers up to ten on chromosome 3. Several genes genes from highly amplified regions of R4 with differential gene expression are indicated in Fig. 8 (SI E)

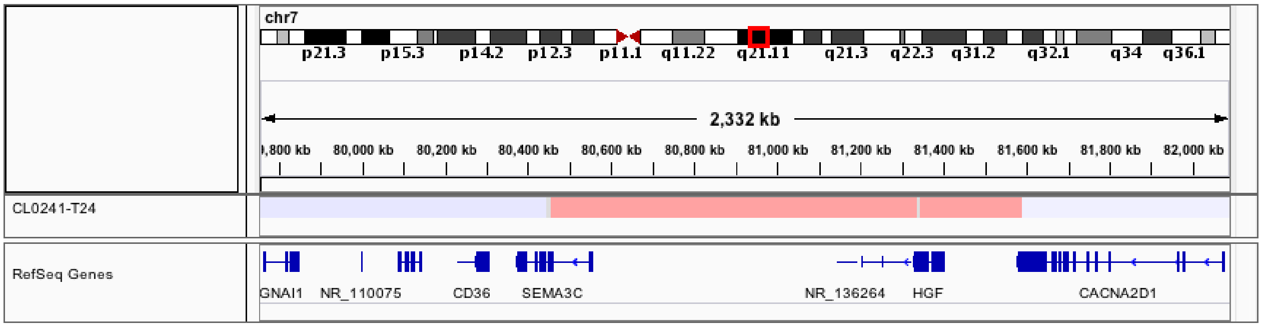


**Figure 7:** Zooming in on the focal *HGF* amplification in R3. The amplicon has a length of approximately 1200 Kb and no other genes than *HGF* lie fully in the amplified region.

### E Gene expression in resistant lines R1–R4

For RNA-seq of PC9 parental and R1-4 derivatives RNA was isolated using the RNeasy Mini Prep Kit (Qiagen, Germany). Library preparation for 3 UTR RNA sequencing was prepared from 500 ng total RNA using the QuantSeq 3 mRNA-Seq library kit (Lexogen, Austria) as described previously [17]. For RNA-seq of EOTD drug-tolerant persisters (EOTD-DTP), PC9 cells were treated for $10-14$ days with EOTD, trypsinized, washed in PBS, and collected by centrifugation. EOTD concentrations were 100 nm erlotinib, 20 nM osimertinib, 10 nM trametinib and 10 nM dasatinib. Due to the low number of EOTD-DTPs, RNA extraction was performed with the Arcturus PicoPure RNA-isolation kit (Thermo Fisher, USA) in 50$\mu$L extraction buffer following the protocol for cell pellets. As comparators, 1000 cells of DMSO treated controls and exEOTD-DTPs were grown without treatment and were extracted in parallel. RNA-seq libraries were prepared using the low-input protocol of the QuantSeq 3mRNA-Seq library kit. RNA-seq libraries were sequenced with a 50bp single-end protocol on an Illumina HiSeq4000 (Illumina, USA). For RNA-seq of Erlotinib DTCs, PC9 cells were grown in 10cm dishes and were treated for 5 days with Erlotinib or DMSO. Total RNA was subsequently extracted using the RNeasy Mini kit (Qiagen, Germany). RNA-seq libraries were prepared from 500ng total RNA with the QuantSeq 3mRNA-Seq library kit standard protocol prior to paired-end sequencing on an Illumina HiSeq4000 (Illumina, USA) with only the first read being used for quantification. Raw sequencing reads were aligned to the human reference genome Hg38 with STAR [18] and gene expression was quantified using RSEM [19] as counts and normalized to library size as counts per million (CPM). Differential gene expression between groups was calculated from count–level data using DESeq2 [20]. The resulting $p$-values were adjusted using the Benjamini-Hochberg correction.

Both R1 and R2 are driven by point mutations in the EGFR pathway. Compatible with that gene expression in those two lines is relatively similar to the parental line, see Fig. 8. In R3, however, we found an amplification of *HGF* which means that the survival signal is now transmitted via a bypass track. This can be seen in the relative gene expression: HGF is highly up-regulated, and also its ligand MET is over-expressed. In R4, HGF and MET expression drop down to the parental level, see Fig. 8 A.

Adding HGF to the parental line abrogates sensitivity to erlotinib, see Fig. 9.

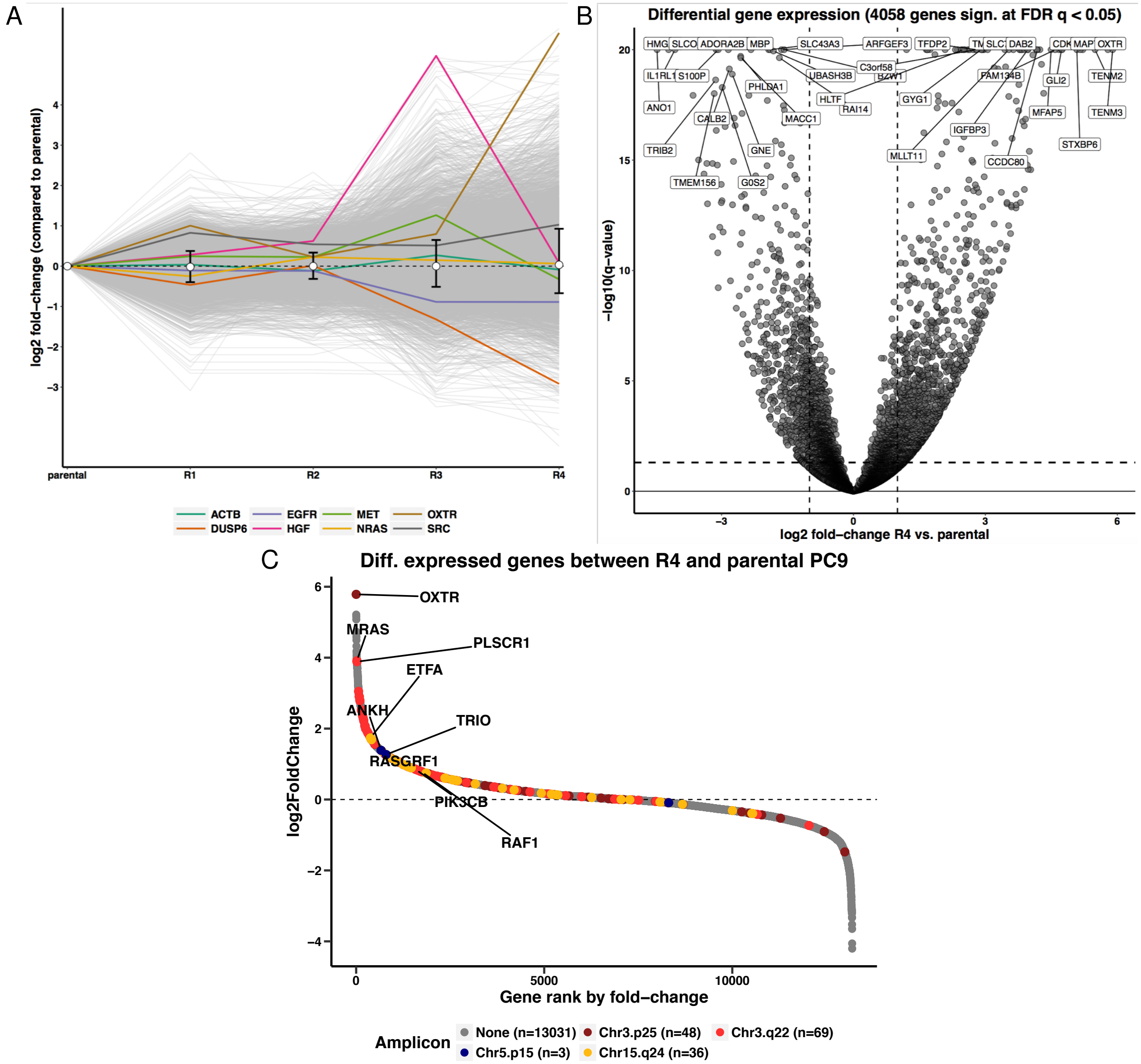


**Figure 8:** A Relative gene expression of the resistant lines as compared to the parental cells. Each line indicates one particular gene. Genes of particular interest are indicated. The difference in gene expression in R3 and R4 compared to the parental line is larger than in R1 and R2. B Genes differentially regulated in R4 relative to the parental line. C Fold changes in R4 vs the gene rank by fold change, with colours indicating the different amplicons found in R4 (see Fig. 6). The number of genes $n$ in each amplicon is indicated in the legend.

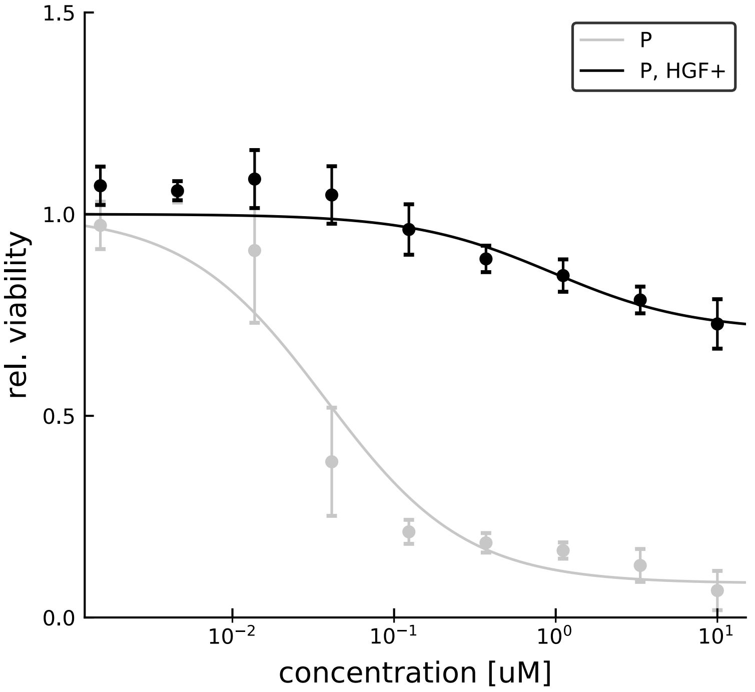


**Figure 9:** HGF concentration and resistance to EGFR-inhibition. Parental cells were treated with 50 ng/ml of HGF. The dose-response curves with and with out HGF show a marked drop in the sensitivity to erlotinib due to the level of HGF.

### F Mutagen treatment

To increase the mutation rate we treated parental cells with N-ethyl-N-nitrosourea (ENU). Cells were seeded and incubated for 24 h under standard growth conditions. The population was then treated with ENU at a concentration of 600 $\mu$g/ml for 30 minutes in serum-free medium under standard cell culture conditions. This treatment increases the point mutation rate approximately by a factor of hundred [21]. The cultures were then washed three times with PBS and cultured for seven days in standard medium (including serum) to allow mutations to fix. After that, the cultures were treated with EOTD to check if colonies grew out.

### G Toxicity tests

To check for toxicity of the EOTD and the EOTDN combinations, we treated cancer cell lines which are not driven by EGFR mutations, specifically HEK293T and Ba/F3, and measured their relative viability (population size after 96 h compared to DMSO control). If the combination were toxic, we expect a decrease of viability of such cell lines. No significant difference in cell viabilities under DMSO and under EOTD treatment is observed for cell lines Ba/F3 (parental and KIF5B-RET transduced) and HEK293T, see Fig. 10A. EOTD concentrations were 100 nm erlotinib, 20 nM osimertinib, 10 nM trametinib and 10 nM dasatinib.

Fig. 10B shows independent experiments comparing viabilities under 100 nm erlotinib, 1 $\mu$M navitoclax, EOTD, and the combination of EOTD and 1 $\mu$M navitoclax (EOTDN). Although Ba/F3 cells show a reduced viability under navitoclax, we find no significant reduction in viability compared to single-drug doses. HEK293T shows no significant reduction viability reduction under EOTDN.

The Ba/F3 cells used in Fig. 10A were a kind gift from Nikolas von Bubnoff. Ba/F3 cells are murine pro-B-cells which are dependent on constitutive IL-3 signaling for survival. Upon transduction with a strong oncogene they become independent of IL-3 signaling but are dependent on the oncogene. Ba/F3 KIF5B-RET cells are dependent on RET signaling and potent RET inhibitors have been shown to kill KIF5B-RET transduced Ba/F3 cells in the low nanomolar range [22].

We also probed the reponse of two further cell lines, H23 and A549 to EOTD and EOTDN, finding a reduced viability under both treatments. Both these cell lines are known to respond already to the combination of trametinib and dasatinib, with a combined (1:1) $IC_{50}$ of 10 nM([23] Fig. 1E, see also [24] Fig. S7) and were thus excluded.

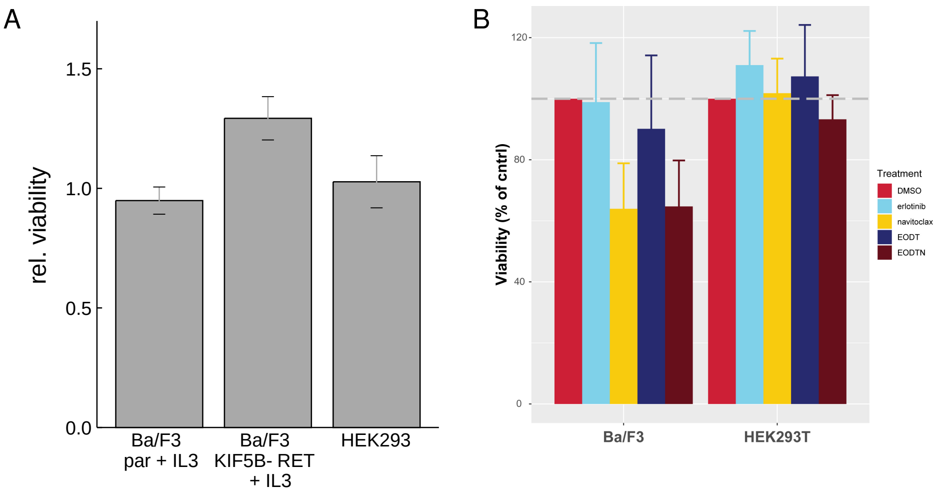


**Figure 10:** Toxicity test of EOTD on different cancer cell lines. A We measured cell viability of EOTD as compared to DMSO control, see text and materials and methods. Experiments were performed in triplicate, the error bars shows the standard error. B Independent experiments, using HEK293T and Ba/F3 cell lines measure cell viability under 100 nm erlotinib, 1 $\mu$M navitoclax, EOTD and EOTDN. Experiments were performed in quadruplicate, the error bars shows the standard error.

### H Drug-tolerant persisters

Drug tolerant cells emerge after the end of prolonged treatment with the EOTD combination, see Fig. 11 for a sample timecourse.

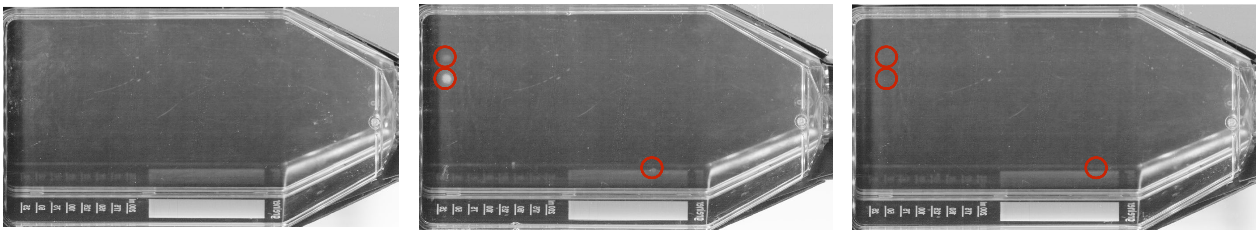

**Figure 11:** PC9 cells under the EOTD combination and after treatment stop. PC9 cells were cultivated in a T175 flask and treated with the EOTD combination. (left) No growing colonies were observed after 7 weeks. At this point, the medium was changed to standard medium. (centre) Three weeks later, three colonies had formed. At this point the medium was switched to 500 nM erlotinib. (right) One week later the colonies had disappeared. We hypothesize that drug tolerant cells had survived the initial EOTD treatment, and some of them switched to a proliferating and erlotinib-sensitive state during the treatment break.

#### H.1 Exiting the drug tolerant state

In principle, the exponential depletion of drug tolerant cells with treatment time (main text Fig. 4) can be explained by two distinct processes. (i) Cells exit the drug tolerant state at a rate $\lambda_{\text{ex}}$ and become sensitive to EOTD treatment. (ii) DTCs are washed out during the medium change. Specifically, the inevitable motion of the medium during medium changes might detach some cells and remove them. To distinguish between these two processes, we vary the frequency at which the medium is changed during our experiments on the rate of DTC depletion.

We plated $10^6$ PC9 cells per flask in T175 culture flasks in regular medium. After one day the EOTD treatment started and was maintained for the prescribed period. Half of the flasks were subjected to two medium changes per week and the remainder was subjected to four medium changes per week. To ensure the DTC at higher rate of medium change are not treated with fresher and thus more potent compounds, we did not add new compounds in these extra changes. Instead, the old medium was centrifuged to separate potentially washed-out cells from the medium, and the medium was returned to the flasks ('sham medium change').

After the prescribed treatment time, the medium was changed back to regular and the cells were left for 1.5 weeks to form colonies that were CV stained and counted. An exponential fit of the form

$$N(t, n) = N_0 \exp\left(-\lambda_{\text{ex}} t - \lambda_{\text{wash}} n\right) \tag{12}$$

to the number of colonies yields the parameters $\lambda_{\text{ex}} = 0.50 \pm 0.03$ week$^{-1}$, $\lambda_{\text{wash}} = 0.031 \pm 0.008$. Here, $n$ is the number of medium changes (both real and sham medium changes). This result indicates that medium change does not contribute significantly to the observed exponential decay of the number of DTC. Figure 12 also shows no significant difference between the depletion of DTC with and without the additional medium changes.

#### H.2 Switching between combination treatment and monotherapy

We look at the response of drug tolerant cells to switching the drug regime from the EOTD treatment to erlotinib monotherapy, and vice versa. Fig. 5A in the main text shows shows typical microscopic images and Fig. 13 shows the corresponding wells stained with CV.

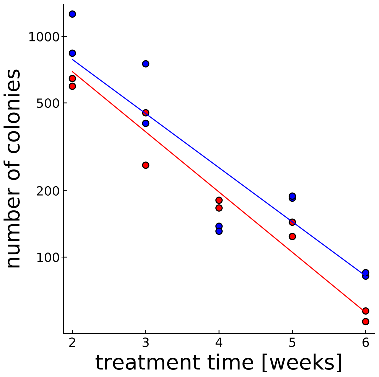


**Figure 12:** The number of colonies formed after treatment with EOTD is plotted against the treatment period indicated (log-scale on the $y$-axis). Blue points indicate the number of colonies for two medium changes per week, red points the number of colonies with additional sham changes, see text. The corresponding lines give exponential fits to the data.

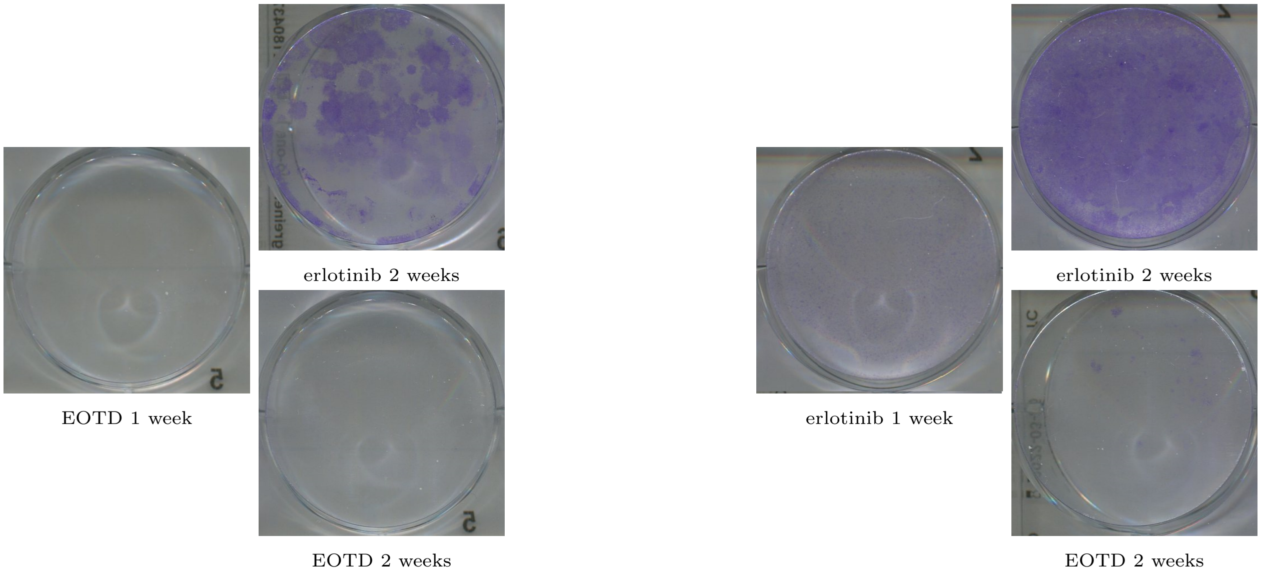


**Figure 13:** The CV stained wells corresponding to the microscopic images shown in Fig.5A of the main text with the same dosing.
